# Longitudinal single-cell epitope and RNA-sequencing reveals the immunological impact of type 1 interferon autoantibodies in critical COVID-19

**DOI:** 10.1101/2021.03.09.434529

**Authors:** Monique G.P. van der Wijst, Sara E. Vazquez, George C. Hartoularos, Paul Bastard, Tianna Grant, Raymund Bueno, David S. Lee, John R. Greenland, Yang Sun, Richard Perez, Anton Ogorodnikov, Alyssa Ward, Sabrina A. Mann, Kara L. Lynch, Cassandra Yun, Diane V. Havlir, Gabriel Chamie, Carina Marquez, Bryan Greenhouse, Michail S. Lionakis, Philip J. Norris, Larry J. Dumont, Kathleen Kelly, Peng Zhang, Qian Zhang, Adrian Gervais, Tom Le Voyer, Alexander Whatley, Yichen Si, Ashley Byrne, Alexis J. Combes, Arjun Arkal Rao, Yun S. Song, UCSF COMET consortium, Gabriela K. Fragiadakis, Kirsten Kangelaris, Carolyn S. Calfee, David J. Erle, Carolyn Hendrickson, Matthew F. Krummel, Prescott G. Woodruff, Charles R. Langelier, Jean-Laurent Casanova, Joseph L. Derisi, Mark S. Anderson, Chun Jimmie Ye

## Abstract

Type I interferon (IFN-I) neutralizing autoantibodies have been found in some critical COVID-19 patients; however, their prevalence and longitudinal dynamics across the disease severity scale, and functional effects on circulating leukocytes remain unknown. Here, in 284 COVID-19 patients, we found IFN-I autoantibodies in 19% of critical, 6% of severe and none of the moderate cases. Longitudinal profiling of over 600,000 peripheral blood mononuclear cells using multiplexed single-cell epitope and transcriptome sequencing from 54 COVID-19 patients, 15 non-COVID-19 patients and 11 non-hospitalized healthy controls, revealed a lack of IFN-I stimulated gene (ISG-I) response in myeloid cells from critical cases, including those producing anti-IFN-I autoantibodies. Moreover, surface protein analysis showed an inverse correlation of the inhibitory receptor LAIR-1 with ISG-I expression response early in the disease course. This aberrant ISG-I response in critical patients with and without IFN-I autoantibodies, supports a unifying model for disease pathogenesis involving ISG-I suppression via convergent mechanisms.

## Introduction

The COVID-19 pandemic has led to the infection of at least 100 million individuals worldwide and over 2.2 million deaths. A perplexing aspect of its pathogenesis is the extreme clinical heterogeneity of infected individuals, with ∼15% of symptomatic patients and < 10% of infected subjects presenting with severe forms of the disease, as defined by dyspnea, pulmonary infiltrates on lung imaging, and low blood oxygen saturation *(1-4)*. Overall, 26.8% of hospitalized patients develop critical disease defined as category 7 on the NIH ordinal scale requiring mechanical ventilation *(5)*. These patients are at the greatest risk for poor outcome and place the most significant burden on the health care system. Despite increasing vaccine availability, some vulnerable individuals may develop critical disease prior to and even perhaps despite vaccination, especially in the context of emerging highly transmissible, more virulent, and antigenically distinct variants of SARS-CoV-2 isolates *(6-11)*. Thus, there is a need to disentangle the immunological consequences of SARS-CoV-2 infection and the underlying immunological causes of critical COVID-19, for stratifying patients early in their disease course and for targeting treatment using available or novel therapies.

Evidence is emerging that genetic and immunological features that pre-date SARS-CoV-2 infection could play an unexpected pathogenic role in severe disease *(12)*. Among patients with critical COVID-19, these features include inborn errors of type I IFN immunity *(13)* as well as the production of autoantibodies against type I interferons (IFNs) *(14, 15)*. Remarkably, these autoantibodies, which seldom occur in healthy controls (< 0.3%) and have not been found in asymptomatically infected subjects, are observed in at least 10% of critical COVID-19 cases *(14)*. The causal relationship between autoantibodies against type I interferons and COVID-19 severity has been supported by their documentation prior to infection and their frequent occurrence in patients with genetic disorders, such as autoimmune polyglandular syndrome type 1 (APS-1) *(15, 16)*.

However, it remains to be determined whether autoantibodies to type I IFNs occur in COVID-19 patients who do not require mechanical ventilation, whether they fluctuate longitudinally during the disease course, and what their consequences are on the composition and phenotypes of circulating leukocyte subsets. Further, few studies have examined circulating leukocytes over the course of SARS-CoV-2 infection *(17, 18)* or have compared with patients presenting with similar respiratory manifestations requiring hospitalization due to other causes *(19)*. Insights into how the natural innate and adaptive immune responses longitudinally evolve in response to SARS-CoV-2 infection, both in anti-type I IFN autoantibody positive and negative cases, may enable the early identification of patients who are likely to develop life-threatening COVID-19 and the discovery of general innate and adaptive mechanisms that can be targeted by therapy.

## Results

### Prevalence of anti-IFN-α2 antibodies in San Francisco

Given the recent description of neutralizing type I IFN autoantibodies in > 10% of critical COVID-19 cases, we sought to determine the frequency of these antibodies in San Francisco in a total of over 4,500 individuals divided over: 1) SARS-CoV-2 positive subjects that span the NIH COVID-19 severity scale *(20)*, 2) a largely asymptomatic community population, and 3) convalescent serum samples from patients previously infected with SARS-CoV-2.

We first determined the frequency of autoantibodies to the type I IFN IFN-α2 in 284 subjects with confirmed SARS-CoV-2 infection using a radioligand binding assay (RLBA). These patients were categorized using the NIH ordinal scale with those scoring between 1-4 classified as moderate, those scoring 5 or 6 as severe, and those scoring 7 as critical (**Table 1**). As positive controls, we also tested 4 COVID-19 negative subjects with APS-1 (**Fig. 1a**). The 284 patients ranged in age from 0-90+ years, were 69% male, had at least one positive SARS-CoV-2 PCR test, and varied in disease severity (**Table 2, 3** and **Table S1, S2**). We found the prevalence of anti-IFN-α2 autoantibodies to be 5/26 (19%) in critical, 6/102 (6%) in severe and absent in moderate disease (**Fig. 1a**). The positive patients were aged 28 to 72 years (mean = 55.7, std = 12.2) and 9/11 (82%) were male. The prevalence of anti-IFN-α2 in critical COVID-19 and the trend of positive patients toward male sex and advanced age is consistent with previously published descriptions *(14)*.

**Table 1:** Categorization of moderate, severe and critical patients in COMET and SFGH cohorts.

| NIH ordinal scale | WHO criteria | categorization: COMET cohort | categorization: SFGH cohort |
| --- | --- | --- | --- |
| 1 | Not hospitalized, no limitations on activities | n/a | moderate |
| 2 | Not hospitalized, limitation on activities and/or requiring home oxygen | n/a | moderate |
| 3 | Hospitalized, not requiring supplemental oxygen – no longer requires ongoing medical care | moderate | moderate |
| 4 | Hospitalized, not requiring supplemental oxygen – requiring ongoing medical care | moderate | moderate |
| 5 | Hospitalized, requiring supplemental oxygen | severe | severe<br>(oxygenation level information not available) |
| 6 | Hospitalized, on non-invasive ventilation or high flow oxygen devices | severe |  |
| 7 | Hospitalized, on mechanical ventilation or ECMO | critical | critical |

**Figure 1:**
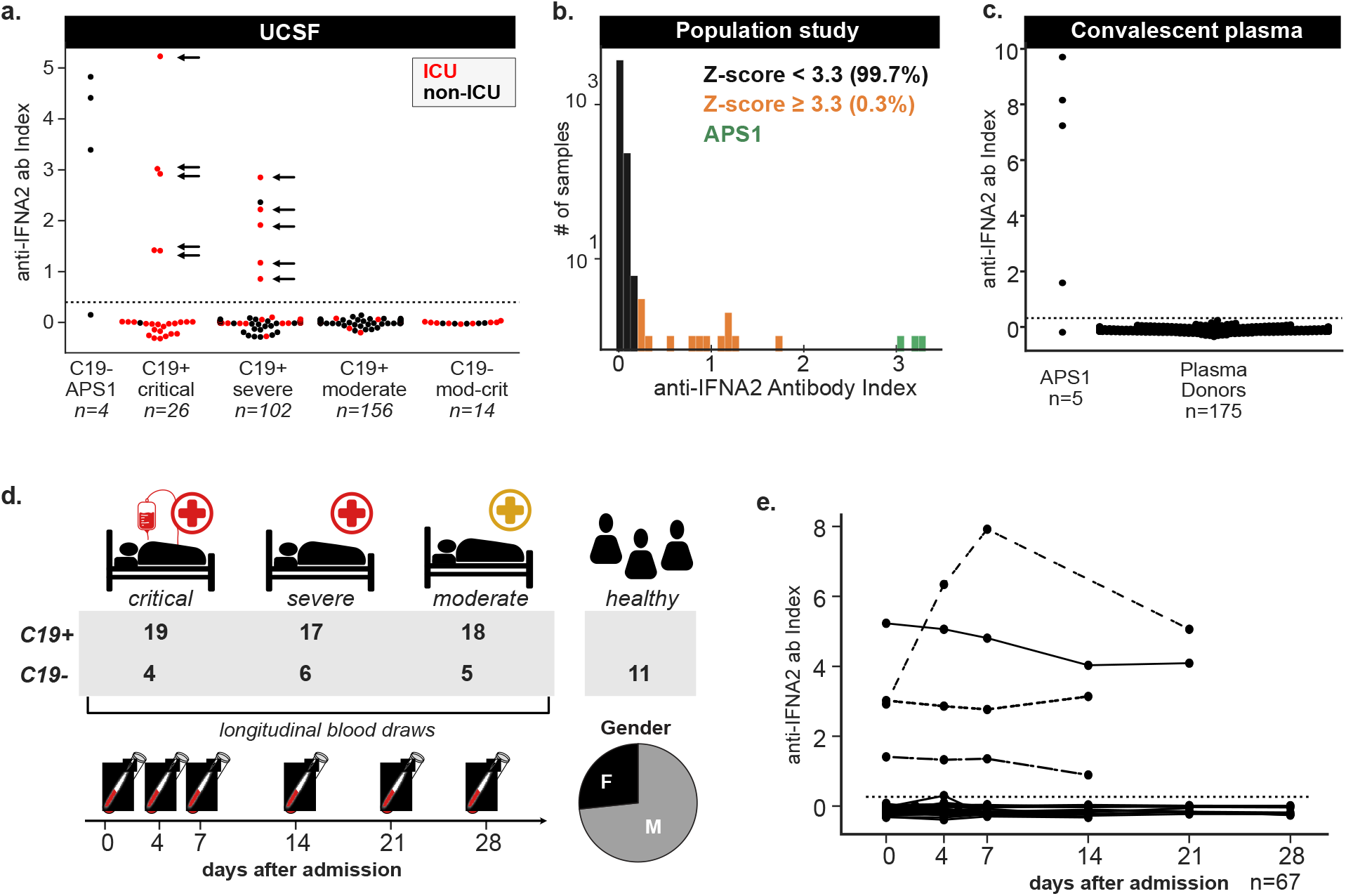
Anti-IFN-α2 antibodies in moderate to critical COVID-19. **a)** Anti-IFN-α2 index (y-axis) in four APS-1 patients, 156 moderate C19+ cases, 102 severe C19+ cases, and 26 critical C19+ cases separated by disease severity and colored by hospitalization status. Positive samples were tested for neutralization against IFN-α2, with arrows indicating those samples with partial or full neutralization ability. Dotted line indicates 6 standard deviations above healthy control mean. **b)** Distribution of the anti-IFN-α2 index across 4,041 subjects in a community cohort from the San Francisco Mission District. **c)** Anti-IFN-α2 index (y-axis) in five additional APS-1 patients and 175 convalescent plasma donors from the Vitalant Blood Center. **d)** COVID-19 Multi-Phenotyping for Effective Therapies (COMET) cohort disease status, severity and gender breakdown. **e)** Anti-IFN-α2 index over days since first hospitalization for 53 hospitalized COVID-19+ and 14 COVID-19- COMET samples. For 2/69 COMET samples anti-IFN-α2 titers were not assessed.

We next examined a community cohort collected during a study of SARS-CoV-2 transmission in San Francisco *(21)* (**Fig. 1b, Table 4, Table S3**). The cohort consists of 4,041 subjects aged 4 to 90 years of Caucasian (36%), Hispanic/LatinX (33%), Asian/Pacific Islander (9%), Black/African American (2%), and other or unknown (20%) descent. In this cohort, a total of 13 anti-IFN-α2 positive individuals (0.32%) were identified. Of these, 5 were male, 6 were female, and 2 were of unknown gender, and positive samples were identified across all represented ethnic groups. None of the participants who were confirmed positive for past or present SARS-CoV-2 infection (117/3,851 by serology and 64/3,758 by PCR) were positive for anti-IFN-α2 antibodies, and all were ambulatory or asymptomatic at the time of testing. These data are consistent with the previously reported absence of autoantibodies in ambulatory COVID-19 patients *(14)*. Our results also confirm the low frequency of anti-IFN-α2 antibodies in individuals independent of, and likely prior to, infection with SARS-CoV-2.

In addition to assessing the presence of anti-IFN-α2 antibodies in San Francisco community cohorts, we analyzed aliquots of convalescent plasma from a central blood bank supplier, encompassing 175 unique plasma donors who had recovered from SARS-CoV-2. Compared with five additional APS-1 subjects, we found that none of the donors tested positive for anti-IFN-α2 autoantibodies (**Fig. 1c**). Reassuringly, this latter cohort suggests that these potentially harmful autoantibodies are rare or absent in the supply from convalescent donors.

### Profiling leukocytes in critical COVID-19 cases with and without anti-IFN-α2 antibodies

We next sought to specifically assess the effects of anti-IFN-α2 autoantibodies in COVID-19 patients on the composition, transcript abundance, and surface protein abundance of circulating leukocytes. For this, we leveraged the COVID-19 Multi-Phenotyping for Effective Therapies (COMET) cohort in San Francisco where peripheral blood mononuclear cells (PBMCs) and serum were longitudinally collected from 69 hospitalized patients presenting with COVID-19 symptoms, of whom 54 were positive (C19+) and 15 were negative (C19-) for SARS-CoV-2, in addition to 11 healthy controls (**Fig. 1d**). Of the C19+ cases, 18 presented with moderate disease, 17 with severe disease, and 19 with critical disease according to the NIH severity scale *(20)* at the time of hospitalization (**Table 1** and **3, Table S2, S5**). For 8/54 C19+ patients the severity changed over the course of hospitalization, of whom 6 improved and 2 worsened. The studied hospitalized patients were ethnically diverse, skewed older than the general population (mean = 59, range = 25 – 90) and were predominantly male (47 men, 22 women) (**Fig. 1d**). While all C19- cases presenting with symptoms concerning for COVID-19 tested negative for SARS-CoV-2, many were infected with common respiratory pathogens confirmed by metagenomic sequencing (**Table S2**). Within the COMET cohort, we identified 4/19 (21%) of the critical COVID-19 cases and none of the moderate to severe cases to be positive for anti-IFN-α2 antibodies (**Fig. 1a**). All 4 cases had anti-IFN-α2 antibodies at the earliest time of sampling and the level of anti-IFN-α2 antibodies remained stable for 3/4 cases across their disease course (**Fig. 1e**).

For immune cell profiling, we collected ∼200 PBMC samples from up to 4 longitudinal timepoints: 0, 4, 7, and 14 days since hospitalization. Multiplexed single-cell epitope and transcriptome sequencing (muxCITE-seq) was performed across 9 pools of genetically distinct samples to simultaneously measure mRNA abundances transcriptome-wide and surface protein abundances of 189 markers from the same cell (**Fig. 2a, Table S4**). A total of 971,550 cell-containing droplets were sequenced and 600,929 cells remained in the final dataset after quality control and removal of doublets, platelets and red blood cells (see **Methods**). Genetic demultiplexing using Freemuxlet resulted in an average of 3,020 cells per sample (**Fig. S1a**).

**Figure 2:**
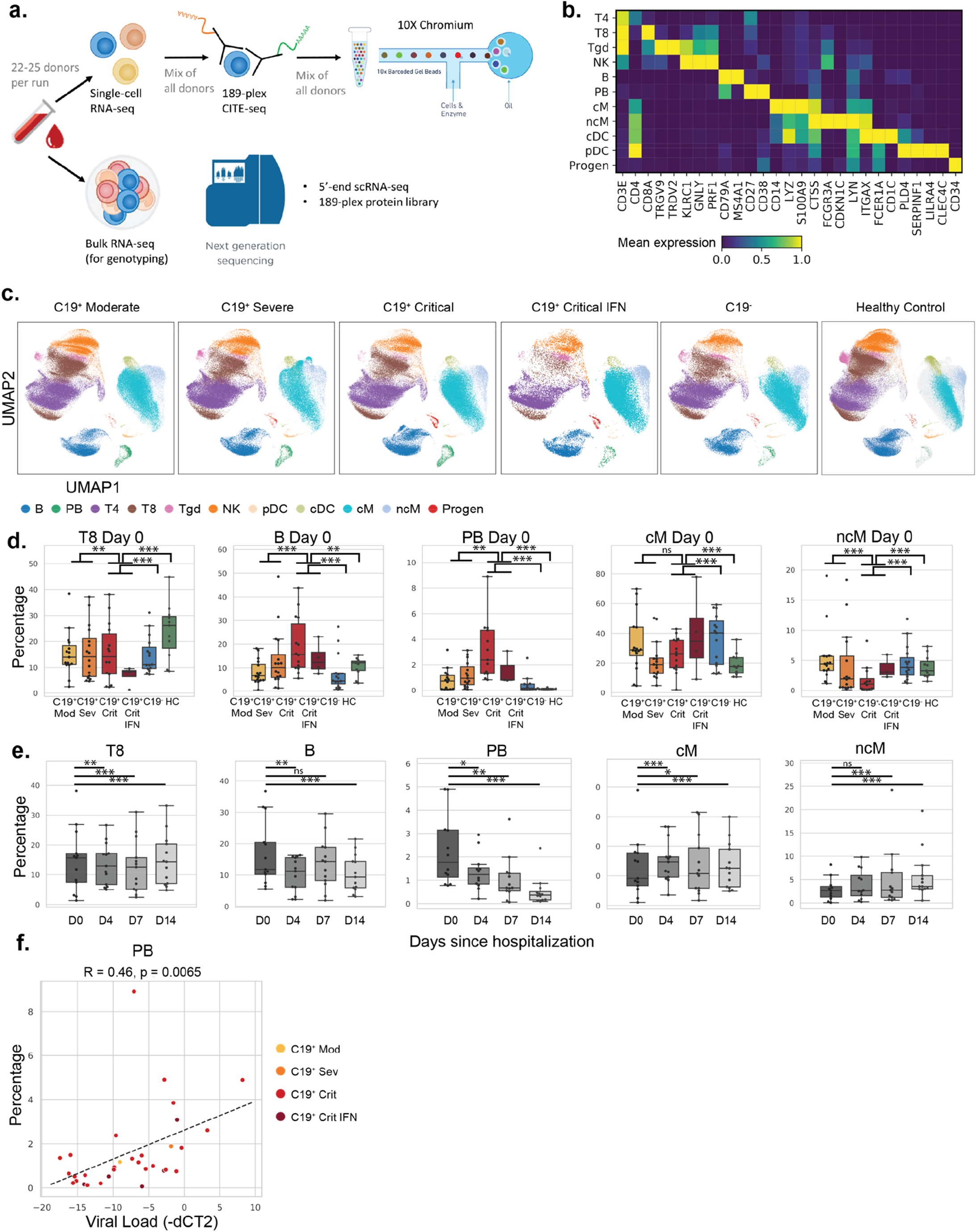
Shifts in circulating leukocyte composition in critical COVID-19. **a)** Experimental set-up. Frozen PBMCs are thawed, multiplexed, stained and processed using cellular indexing of transcriptomes and epitopes by sequencing (CITE-seq). **b)** Marker genes for each of the 11 cell types identified including CD4+ T cells (T4), CD8+ T cells (T8), gamma delta T cells (Tgd), natural killer cells (NK), B cells (B), plasmablasts (PB), classical monocytes (cM), non-classical monocytes (ncM), conventional dendritic cells (cDC), plasmacytoid dendritic cells (pDC), and CD34+ hematopoietic progeneitors (Progen). **c)** UMAP projections of PBMCs from donors separated by COVID-19 status and severity. Cells are colored by type. **d)** Boxplots (showing median, 25^th^ and 75^th^ percentile) of the percentages of T8, PB, cMs and ncMs (y-axis) by COVID-19 status and severity level on day of hospital admission (D0). Each dot represents the percentage of a specific cell type per donor. Shown statistical comparisons are between cells from all C19+ critical donors (including the anti-IFN-α2 autoantibody donors) and healthy controls, C19- donors or combined C19+ Moderate-Severe donors. Other cell types can be found in **Fig. S1. e)** Boxplots of the percentages of T8, PB, cMs, ncMs (y-axis) in COVID-19 patients over day 0, 4, 7 and 14 since hospitalization (D0, D4, D7, D14). Other cell types can be found in **Fig. S1. f)** Scatterplot of SARS-CoV2 viral titer as measured by qRT-PCR in tracheal aspirates (inverse dCT, x-axis) and percentage of plasmablasts (PB) (y-axis) as quantified in donor-matched single-cell PBMC data (R = Pearson correlation). Holm’s multiple-testing corrected, permutation-based p-values: *** p < 0.001, ** p < 0.01, * p < 0.05, ns = not significant.

### Critical COVID-19 is characterized by increased frequency of plasmablasts and classical monocytes

We compared the frequencies of 11 cell types defined using a combination of mRNA and surface protein markers between C19+ cases, C19- cases, and controls, as well as within C19+ cases separated by severity (see **Methods**). The assessed cell types include plasmablasts (PB), B cells (B), CD4^+^, CD8^+^, and gamma delta T cells (T4, T8, Tgd), natural killer cells (NKs), conventional and plasmacytoid dendritic cells (cDC and pDC), classical and non-classical monocytes (cM and ncM), and hematopoietic progenitor cells (Progens) (**Fig. 2b**). We first confirmed that muxCITE-seq-derived estimates of lymphocyte and monocyte frequencies were well correlated with complete blood count measurements reported in the electronic health record from the same donor within +/- 2 days (Pearson R_monocyte_= 0.59, P = 7.2×10^−105^; Pearson R_lymphocyte_ = 0.57, P = 8.2×10^−55^; **Fig. S1b**). Qualitatively, C19+ cases exhibited shifts in the Uniform Manifold Approximation and Projection (UMAP) space of circulating leukocytes, particularly of myeloid cells, that were not confounded by processing batch and pool (**Fig. 2c, Fig. S1c**). Comparing critical C19+ cases to healthy controls, we observed statistically significant changes in frequencies for every cell type, including prominent increases in the frequencies of B, PB and cMs (cM: median change +10.0%, Differential proportion analysis (DPA) permutation P < 10^−5^; B: +2.7%, P = 2.1×10^−3^; PB: +2.1%, P < 10^−5^), and decreases in the frequencies of T8 and Tgd (T8: -15.4%, P < 10^−5^; Tgd: -3.9%, P < 10^−5^). These changes in T8, Tgd and PB were most significant in critical C19+ cases and the frequencies in moderate and severe cases were between those observed in critical cases and healthy controls (**Fig. 2d, Fig. S1d, Table 5, Methods**). Interestingly, the frequency of T8s were even lower and the frequency of cMs were even higher in critical C19+ cases with detectable anti-IFN-α2 antibodies than those without (T8: -5.8%, P < 10^−5^; cMs: +8.3%, P = 0.034) (**Fig. 2d, Fig. S1d, Table 5**). Importantly, the described changes in frequencies were significantly different between critical C19+ cases and C19- hospitalized patients, suggesting these effects are not likely explained by hospitalization in general (**Fig. 2d, Fig. S1d, Table 5**). For the 14 C19+ donors for whom all 4 timepoints were available, we observed decreases in the frequencies of B and PB cells over time (Median change D0 vs D14; B: -3.7%, P = 6.0×10^−5^; PB -0.8%, P < 10^−5^) and increases in the frequencies of cM and ncMs (D0 vs D14: cM +5.7%, P < 10^−5^; ncM +2.5%, P < 10^−5^) for both days since hospitalization and days since onset of first symptoms (**Fig. 2e, Fig. S1e, S1f, Table 5, Methods**). These longitudinal changes normalized towards frequencies observed in healthy controls, except for the frequency of cMs which appears to further increase from levels observed in healthy controls. Previously, the frequency of PBs has been observed to correlate with COVID-19 disease severity *(22)* and to diminish upon recovery *(18)*. We observed that the reduced PB frequency was positively correlated with reduced viral titer over time (Pearson R = 0.46, P_adjusted_ = 0.0065) suggesting coordinated dynamic changes of host humoral immunity and viral load over the course of hospitalization (**Fig. 2f, Fig. S1g, S1h**). Overall, these analyses revealed shifts in cell type composition specific to COVID-19, between patients of varying disease severity, and over time. The general comparable composition of circulating leukocytes in critical C19+ patients with and without anti-IFN autoantibodies suggests the presence of a broader, conserved mechanism underlying severe disease, such as additional IFN-related pathology particularly in the autoantibody negative patients.

### Critical COVID-19 is marked by deficient type-1 ISG expression early in disease course

To further characterize cell-type intrinsic changes in COVID-19 and assess the effects of anti-IFN-α2 antibodies, we compared mRNA and surface protein abundances between C19+ cases, C19- cases, and healthy controls for each cell type. We identified 161 genes (FDR < 0.05, log2(fold change) log_2_FC > 1) whose transcripts were differentially upregulated between C19+ cases at day 0 and healthy controls in at least one cell type (**Fig. 3a, Fig. S2a, Table 6**). K-means clustering of the 161 differentially expressed genes aggregated for each of 11 cell types at day 0 identified five clusters, including a cluster (cluster 1) of genes enriched for type I IFN signaling and viral response primarily expressed in myeloid cells (GSEA: Type I IFN signaling pathway, permutation P < 10^−5^), a cluster (cluster 2) enriched for neutrophil degranulation (GSEA: neutrophil degranulation, P < 10^−5^), a cluster (cluster 3) of immunoglobulins and plasmablast activation markers, and a cluster (cluster 4) enriched for complement activation in non-classical monocytes (GSEA: complement activation, P = 0.026) (**Table S6**). Given the heterogeneous expression of the IFN signaling cluster (cluster 1) within COVID-19 patients, we further compared the expression of type I-specific and type II-specific ISGs between C19+ cases and healthy controls and within C19+ cases of varying severity. To differentiate type I- and II-specific ISGs, we compared healthy donor PBMCs stimulated with recombinant IFN-beta or IFN-gamma from an independent single-cell RNA-sequencing (scRNA-seq) dataset to identify genes specifically upregulated by either interferon (**Fig. 3b**, see **Methods**). Strikingly, in myeloid cells (cM, ncM, pDC, cDC), the average expression of type I-specific and to a lesser extent type II-specific ISGs in critical cases on day 0 of hospitalization was significantly lower compared to moderate and severe cases (type I: log_2_FC = -0.51 to -0.82, P = 2.2×10^−4^ to 1.7×10^−3^; type II, cM only: log_2_FC = -0.46, P = 7.6×10^−4^) (**Fig. 3c, Fig. S2b, S2c**). We also found that the expression of type I-specific ISGs in the four critical C19+ cases with anti-IFN-α2 autoantibodies was the lowest among the C19+ cases at levels observed in healthy controls (C19+ critical IFN vs healthy: n.s. in cM, ncM, cDC, pDC) (**Fig. 3c**). Through the disease course, average expression of type I-specific ISGs in moderate and severe cases was high at the time of hospitalization but quickly diminished, while in critical cases, especially those with anti-IFN-α2 autoantibodies, average expression of type I-specific ISGs remained low (**Fig. 3d**). These findings suggest that there is a shared causal mechanism of critical disease in patients with and without autoantibodies to type I IFNs. The latter patients may have undetected or other acquired or inherited defects in the type I IFN immune response.

**Figure 3:**
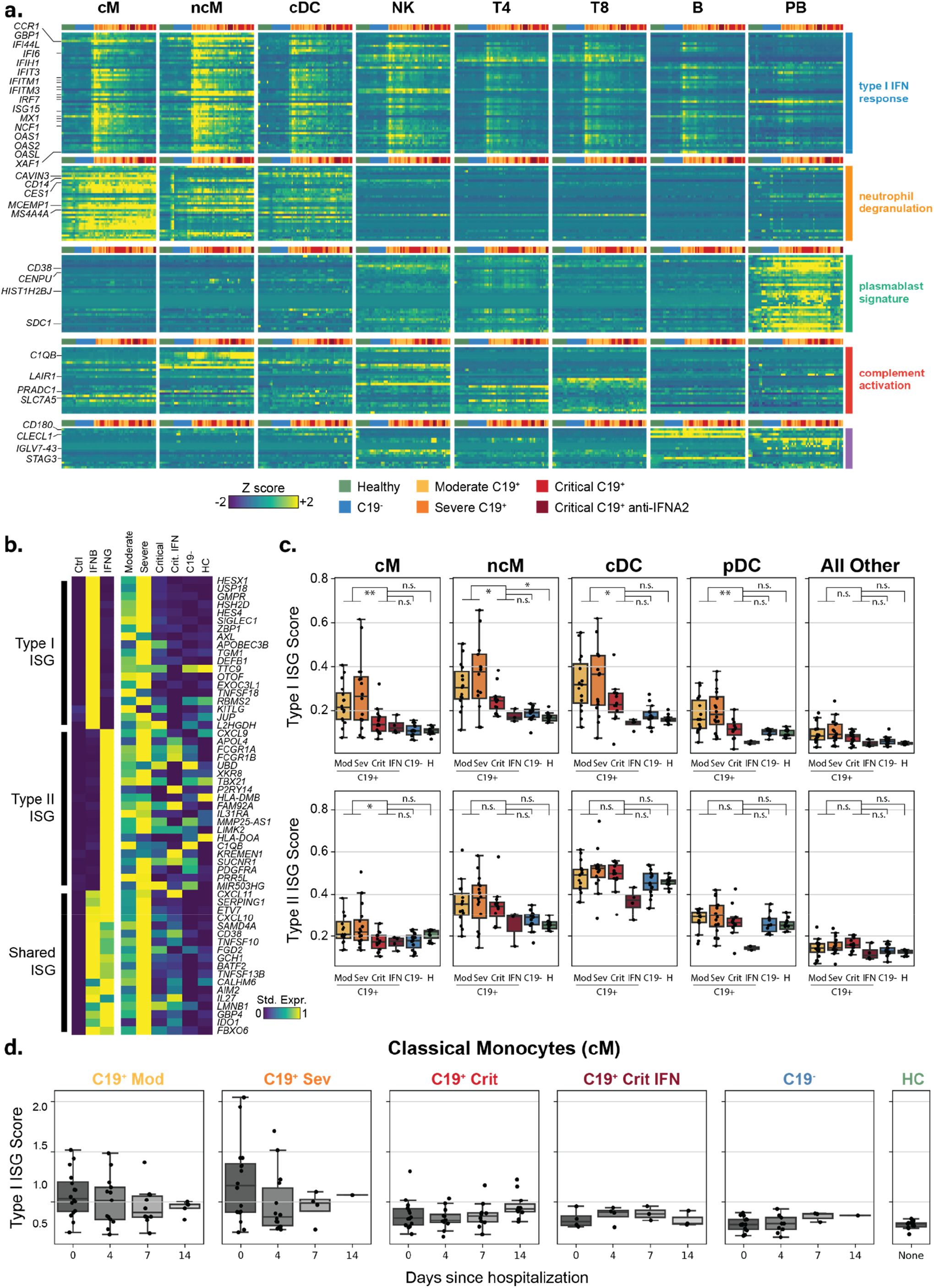
Transcript abundance changes of leukocyte subsets in critical COVID19. **a)** Heatmap of 161 differentially expressed genes at day 0 (FDR < 0.01, |log(fold change)| >1) in at least one of 11 cell types. CD4+ T cells (T4), CD8+ T cells (T8), natural killer cells (NK), B cells (B), plasmablasts (PB), classical monocytes (cM), non-classical monocytes (ncM), and conventional dendritic cells (cDC) are shown. Each row represents a gene and each column is the average expression of the genes in a particular sample across all cells of a specific type. Samples are grouped by both cases control status and C19+ severity. Expression levels are row standardized. Genes are grouped by cluster with the enriched clusters annotated. **b)** Matrix plot of type I and type II-specific ISGs defined using an orthogonal scRNA-seq data set (left plot) and in the COMET cohort separated by case control status and disease severity (right). **c)** Type I and type II-specific ISG scores (y-axis) at day 0 across 4 myeloid cell types, and pseudobulk of all other cell types, separated by case control status and disease severity. Boxplots show median, 25^th^ and 75^th^ percentile. Cell types comprising the pseudobulk are in supplementary materials. **d)** Type I-specific ISG score (y-axis) over the course of disease for healthy controls, C19- and C19+ cases in classical monocytes. C19+ cases are separated by severity and the presence of anti-IFN-α2 antibodies. *** p < 0.001, ** p < 0.01, * p < 0.05, ns = not significant.

### Type I ISG deficiency is inversely correlated with surface expression of LAIR-1

We next sought to identify changes in the expression of surface proteins in COVID-19 patients that may be correlated with type I ISG expression. The correlation of surface protein and transcript abundance varied across the 189 targeted genes with lineage specific surface markers exhibiting the highest correlation (**Fig. S3a**). Comparing C19+ cases at day 0 of hospitalization to healthy controls for each cell type separately, we identified 5/189 differentially expressed surface proteins in cMs and an additional 14/189 in other cell types (**Fig. 4a, Fig. S3b**, |log_2_FC| > 0.5, FDR < 0.05). Of the five proteins differentially expressed in cMs, four (TFRC, SIGLEC-1, FCGR1A, and LAIR1) were higher expressed in C19+ cases. SIGLEC-1 is a known up-regulated ISG whose pattern of surface expression is consistent with the expression of type I-specific ISGs (**Fig. 3c, Fig. 4b**). In cMs but not other cell types, leukocyte-associated immunoglobulin-like receptor 1 (LAIR-1) was differentially up-regulated in critical C19+ cases compared with healthy controls (log_2_FC = 0.88, p < 9.8×10^−6^) and moderate/severe C19+ cases (log_2_FC = 0.47, p < 2.0×10^−3^) (**Fig. 4b, Fig. S3b**). Further, the four critical C19+ samples with anti-IFN-α2 autoantibodies were among the top 10 with the highest LAIR1 expression in cMs. Note that although LAIR-1 was an inhibitory molecule discovered in lymphocytes, its expression and differential expression between cases and controls were highly specific to cM and ncM cells and not to T4, T8, NK, B or PBs (**Fig. 4b**). Interestingly, LAIR-1 expression in cMs from critical C19+ cases was high early in the disease course and diminished over time, inversely tracking with the pattern observed for type I-specific ISGs (**Fig. 4c, Fig. S3c**). Further, in C19+ cases at day 0, the expression of surface LAIR-1 was inversely correlated with expression of type I-specific ISGs in cMs (Pearson R = -0.47, p < 0.01) and ncMs (Pearson R = -0.41, p < 0.01) (**Fig. 4d, 4e, Fig. S3d**). Unlike down-regulated ISG surface proteins (e.g. CD244, SLC3A2) that were also inversely correlated with the type I-specific ISG score (**Fig. 4d**), LAIR-1 is not expressed in healthy samples suggesting that it is not a down-regulated ISG. These results demonstrate LAIR-1 as a highly specific monocyte cell-surface biomarker predictive of deficient type I-specific interferon response.

**Figure 4:**
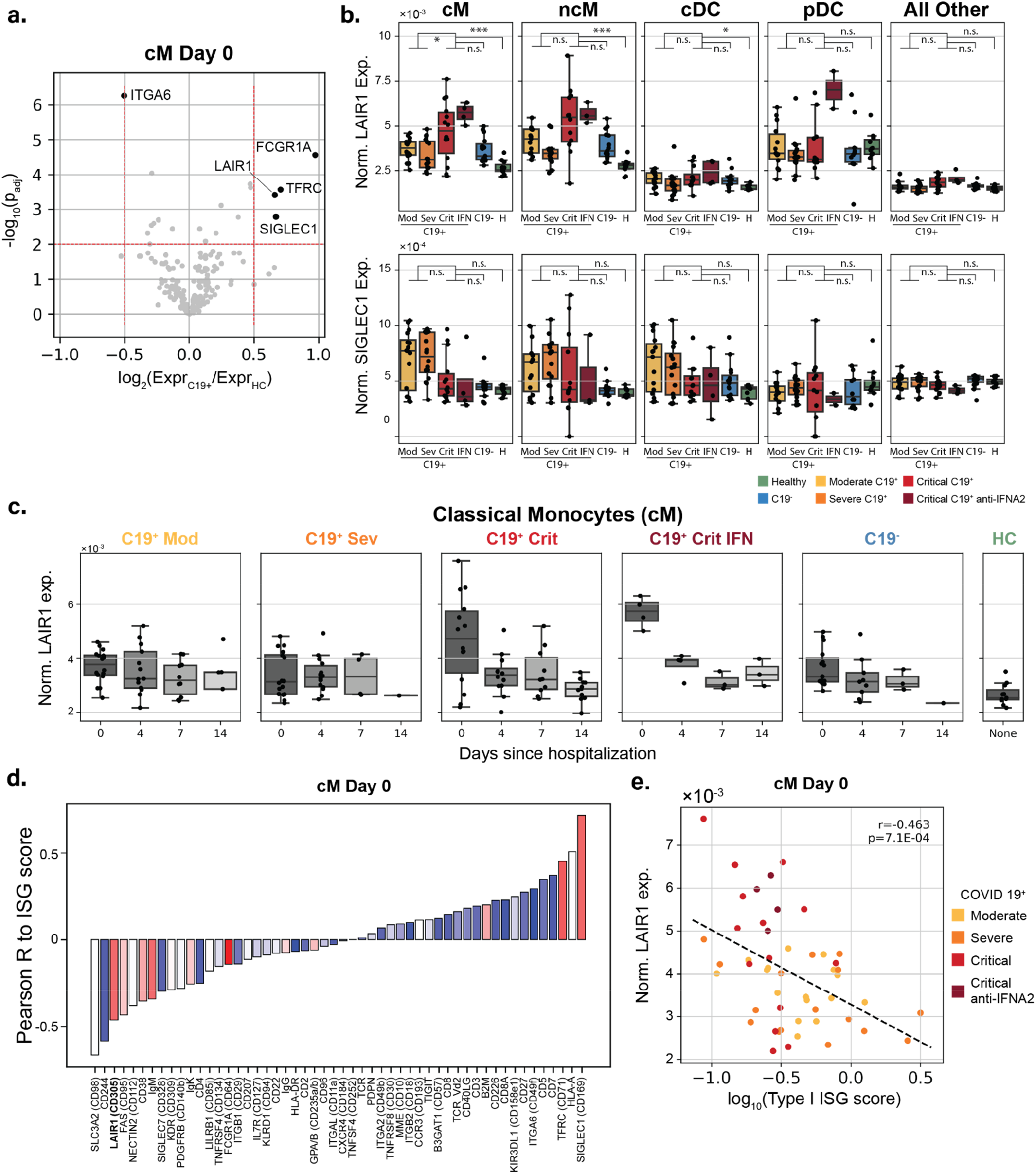
Surface protein abundance changes of leukocyte subsets in critical COVID19. **a)** Volcano plot of log fold change between C19+ and healthy controls (x-axis) versus -log10(P-value) (y-axis) in cM cells. Proteins that are statistically significant (FDR < 0.05) and have a log2(fold change) > 0.5 are highlighted. **b)** Normalized LAIR-1 and SIGLEC-1 surface expression (y-axis) at day 0 across 8 cell types separated by case control status, severity, and presence of anti-IFN-α2 antibodies. Boxplots show median, 25^th^ and 75^th^ percentile. Tgd, pDC, Progens are in **Fig. S3. c)** Normalized LAIR-1 surface expression (y-axis) in classical monocytes over the course of disease for healthy controls, C19- controls, and C19+ cases. C19- controls and C19+ cases are separate by severity and the presence of anti-IFN-α2 antibodies. **d)** Barplot of correlation between the surface expression of xx statistically significantly proteins to type I-specific ISG score in classical monocytes at day 0. Proteins are colored by their log2(fold change) expression between C19+ cases and healthy controls. Red: Higher expression in C19+ cases. Blue: Lower expression in C19+ cases. **e)** Scatterplot of normalized LAIR-1 expression (y-axis) versus the type I-specific ISG score (x-axis) for C19+ cases colored by severity and anti-IFN-α2 status. *** p < 0.001, ** p < 0.01, * p < 0.05, ns = not significant.

## Discussion

The dramatic clinical heterogeneity over the course of SARS-CoV2 infection, ranging from asymptomatic to lethal, is a key observation and defining feature of this pandemic. It is important to understand what causes life-threatening COVID-19 pneumonia in a minority of infected individuals. Recent work has suggested that pre-existing autoimmunity against type I IFN can underlie critical COVID-19 pneumonia in > 10% of the cases *(14)*. Here, we have confirmed that neutralizing autoantibodies to type I IFNs indeed are present in severe to critical COVID-19 patients from two independent cohorts, showing a combined prevalence of about 9%. Both the C19+ and asymptomatic cohorts studied here had significant Hispanic representation, a population that has not been previously studied at scale. The presence of anti-type I IFN autoantibodies in this population indicates that this phenomenon may be widely conserved across a diversity of ancestries. In terms of age and gender, the majority of autoantibody-positive severe to critical COVID cases were male and > 55 years of age, consistent with previous reports; however, these numbers did not reach significance as compared to the observed frequencies among all C19+ patients *(14)*. Interestingly, we observed roughly equal numbers of positive males and females in our community survey. Further study will be required to determine if there exist differences between the autoantibodies in male versus female COVID-naïve patients that could partially explain the downstream skewing of hospitalized patients towards male gender, such as differential neutralization ability or additional accompanying risk factors.

In anti-type I IFN positive critical patients in our longitudinally sampled cohort, we were also able to determine that these autoantibodies were present from the earliest timepoint in their clinical course (collected within 4 to 13 days from the start of their first disease symptoms). Given the time required for a detectable, stable humoral immune response to form (2-3 weeks) *(23)*, our data strongly suggest that autoantibodies pre-date infection with SARS-CoV2. Consistent with this, our survey of a community cohort in the San Francisco Mission District also revealed a subset of presumed COVID-19- naïve individuals who were anti-type I IFN antibody positive (0.3%), suggesting that there are individuals who may be at higher risk for critical disease due to these pre-existing autoantibodies, including both males and females across a broad range of ages. Moreover, in a community-based population study, we did not detect these autoantibodies in 154 patients with asymptomatic or ambulatory infection with SARS-CoV-2 (compared to 13/3821 uninfected donors) or in convalescent plasma donor samples, confirming that the penetrance of severe to critical COVID-19 in infected individuals with autoantibodies is so far complete.

In addition to validating the presence of anti-type I IFN autoantibodies in severe and critical COVID-19 patients, we further have shown using scRNA-seq that these antibodies are associated with impaired type I ISG response in several distinct myeloid populations. While other similar high-dimensional immune profiling studies have found evidence of impaired ISG responses in monocytes *(24, 25)* and neutrophils *(19)*, we have now provided additional specificity and a clear mechanism of how this may unfold in a subset of subjects. Interestingly, we also find impaired myeloid type I ISG expression in additional critical subjects without detectable anti-type I IFN autoantibodies. This important observation suggests that impaired type I IFN immunity is a shared mechanism of more severe forms of the disease in patients with and without autoantibodies to type I IFNs *(13)*. Patients without detectable autoantibodies may have lower titers of autoantibodies, autoantibodies that neutralize lower amounts of type I IFNs, or autoantibodies undetectable because they are bound to type I IFNs. Alternatively, these patients may carry inborn errors of the production and amplification of type I IFNs, as recently shown in other patients *(13)*, or antibody-mediated mechanisms may exist that are independent of the direct binding to IFNs *(19)*. Genetic and immunological studies are underway in our cohort of patients. These findings, along with the observation of high type I ISG expression in mild patients early during the disease course that quickly diminishes, further suggest that impaired type I IFN immunity during the first hours and days of infection may account for the protracted disease course including pulmonary and systemic inflammation. A two-step model of life-threatening COVID-19 is emerging, with defective type I IFN intrinsic immunity in the first days of infection resulting in viral spread, in turn unleashing leukocyte-mediated excessive inflammation in the lungs and other infected organs during the second week of infection *(12)*.

Our analysis of 189 cell-surface proteins by CITE-seq identified the expression of LAIR-1 in cMs to be elevated in COVID-19 patients and correlated with the impaired type I ISG response. LAIR-1 is an inhibitory surface protein originally discovered in T and NK cells, and is involved in inhibiting NK-mediated cell lysis and effector T cell cytotoxicity upon FcR-mediated cross-linking *(26-28)*. More recently, it has also been shown in monocytes and pDCs that cross-linking of LAIR-1 can inhibit the production of IFNα in response to TLR ligands in healthy controls and patients with systemic lupus erythematosus *(29, 30)*. Importantly, LAIR-1 expression is highest in cMs at the time of initial hospitalization and decreases rapidly by day 4 among a subset of critical patients, including the four with anti-type I IFN autoantibodies. Whether LAIR-1 plays a causal role in deficient type I IFN response would require further investigation. Nevertheless, the ability to use a highly cell-type specific surface protein to predict impaired type I IFN response in critical COVID-19 patients early during disease provides an important biomarker.

Our findings have several important implications for the ongoing pandemic and our understanding of patients with a critical COVID-19 clinical course. First, our results show that an impaired type I ISG response early in the disease course in multiple immune populations is associated tightly with autoantibodies to type 1 interferons, providing a glimpse into the immune dysregulation present in patients with a severe clinical course. In this regard, it is critical to be able to identify patients with an impaired type I ISG response early during disease course; a combination of the highly specific assays for autoantibodies against type I IFNs and biomarkers for deficient ISGs such as LAIR-1 could quickly allow triaging of patients during initial hospitalization. Second, treatment strategies with IFNβ might be particularly valuable for those with preexisting antibodies to type I IFNs. The large immunological differences of severe patients in the earliest timepoints additionally suggest that identification and treatment would likely need to happen early in the disease course. Third, we found that autoantibodies to type I IFNs in severe COVID-19 subjects were present at the time of their presentation and precede the development of antibodies to SARS-CoV2. This, along with the presence of healthy autoantibody-positive individuals in the community, suggests that anti-type I IFN autoantibodies pre-date infection and that there exists an at-risk group for severe disease in the general population. Going forward, strategic efforts to identify this high-risk population early in the disease course could have significant impact on improving clinical outcomes including mortality rates, and identifying these individuals before infection could have a major impact on preventive measures.

**Table 2: Demographics of the SFGH cohort**

Demographics and clinical characteristics of patients from the SFGH cohort, including comparison across anti-INF-α2 positive and negative patients. Significance values were determined using Fisher’s exact test, except in the case of continuous distributions (Length of Stay), where a Kolmogorov-Smirnov test was used.

**Table 3: Demographics of the COMET cohort**

Demographics and clinical characteristics of patients from the COMET cohort, broken down by anti-INF-α2 positive C19+, anti-IFN-α2 negative C19+, and C19- patients.

**Table 4: Demographics of the community cohort**

Demographics of patients from the community cohort, broken down by anti-INF-α2 positive and anti-IFN-α2 negative individuals. Significance values were determined using Fisher’s exact test.

**Table 5: Differential proportion analyses**.

Complete differential proportion analyses results corresponding to **Fig. 2d** (C19+, C19- and healthy controls at time of hospital admission, D0) and **Fig. 2e** (C19+ cases along their hospitalization course, D0-4-7-14). Analyses were performed on the cell observations per disease severity level (cells from all C19+ moderate and C19+ severe donors, C19+ critical donors including the anti-IFN-α2 autoantibody donors, all C19- donors, and healthy controls are combined in 4 separate groups) and per cell type (cellcounts tab). The resulting statistics are based on 100,000 permutations and Holm’s multiple-testing correction (stats tab). Median counts over all donors per disease severity level and cell type are provided in the mediancounts tab.

**Table 6: Differentially expressed genes**.

List of differentially expressed (DE) and variable (DV) genes comparing C19+ vs healthy controls at 0, 4, 7 and 14 days after hospital admission (D0, D4, D7, D14) in each of the 11 defined immune populations.

## Materials and Methods

### Cohorts and patient enrollment

The COVID-19 Multi-Phenotyping for Effective Therapies (COMET) cohort includes patients recruited to the Immunophenotyping Assessment in a COVID-19 Cohort (IMPACC) study. All patients hospitalized for symptomatic COVID-19 infection at both the tertiary care center and the safety-net county hospital associated with the University of California San Francisco, were eligible to participate in the COMET cohort study. Biospecimens may be collected under an IRB-approved initial waiver of consent with subsequent attempts to consent surrogates and study subjects for full study participation. We selected samples from 69/101 of subjects enrolled in COMET between 4/8/2020 and 6/20/2020. Sample selection was prioritized in the patients that were hospitalized with longitudinal blood collections and therefore PBMCs were available during a 14-day time period. This study is approved by the Institutional Review board: UCSF Human Research Protection Program (HRPP) IRB# 20-30497.

Details of the community-based cohort are described in Chamie et al. 2020 *(21)*. APS1 patients in the study were collected at the NIH under Protocol #11-I-0187 and were previously published in Ferre et al. (2016) and Ferre et al. (2019) *(31, 32)*.

Convalescent plasmas (CCPs) were collected in the Vitalant system following FDA Guidance for donor eligibility. These criteria evolved throughout the study period due to testing availability and evolution of the pandemic in the United States. Evidence of COVID-19 was required in the form of a documented positive SARS-CoV-2 molecular or serologic test, and complete resolution of symptoms initially at least 14 days prior to donation but then a minimum of 28 days was implemented. All CCP donors were also required to meet traditional allogeneic blood donor criteria. At the time of plasma collection, donors consented to use of de-identified donor information and test results for research purposes. All CCPs were tested for SARS-CoV-2 total Ig antibody using the Ortho VITROS CoV2T assay at our central testing laboratory (Creative Testing Solutions [CTS], Scottsdale, AZ). CCP qualification requires the signal-to-cutoff ratio S/CO of this test to be at least 1.0. Retention samples of serum and plasma for all donations are archived at the Vitalant Research Institute Denver. Plasma samples are from 175 unique CCP donors where some had repeated donations for a total of 281 samples. These samples were selected solely on the Ortho VITROS CoV2T assay results to represent the entire range of high to low signal to cutoff (S/CO) signal. Collections were from across the US Vitalant system from April 8 to September 1, 2020.

### Isolation and preparation of PBMCs for scRNA-seq

Whole blood from 80 donors was drawn into plastic EDTA Vacutainer blood collection tubes (Becton, Dickinson and Company) at the time of hospital admission (D0) and 4 (D4), 7 (D7) and 14 (D14) days later. Of these donors, 69 were patients with high clinical suspicion of COVID-19 infection that were admitted at UCSF or ZSFG and 11 were healthy donors. COVID-19 status was assessed for all 69 patients by reverse transcriptase polymerase chain reaction (RT-PCR) tests of nasal swab samples and confirmed that 15 patients were COVID-19 negative, whereas 54 patients were COVID-19 positive. During the hospitalization, the severity of each COVID-19 positive patient was assessed using the NIH COVID-19 severity scale (**Table 1**) *(20)*. For all analyses we categorized patients based on their severity at time of hospital admission (D0).

Peripheral blood mononuclear cells (PBMCs) were isolated at RT using SepMate PBMC Isolation Tubes (STEMCELL Technologies) by the UCSF Biospecimen Resource Program. In brief, 7.5mL of whole blood was centrifuged at 1,000 rcf for 10 min with swinging-out rotor and brake off to separate 3.5mL of plasma. Remaining blood was diluted with 8mL of DPBS and slowly added to a SepMate-50 tube pre-filled with 15mL of Lymphoprep (STEMCELL Technologies). The tube was then centrifuged at 800 rcf for 20 min with brakes off. After centrifugation, the top layer including the buffy coat was gently and quickly poured into a 50 mL falcon tube to centrifuge at 400 rcf for 10 min with brake on. The pellet was washed twice each time with 20 mL of EasySep buffer (STEMCELL Technologies) followed by centrifugation at 400 rcf for 10 min with brakes on. Washed PBMCs were counted and resuspended at 5×10^6^ cells/mL in cold freezing media (FBS with 10% DMSO). Cells were aliquoted into cryovials at 1-5×10^6^ cells per vial and transferred in a Mr. Frosty to the -80°C freezer for 24 hours before cryopreservation in liquid nitrogen.

### Single-cell multimodal immunophenotyping

Multiplexed single-cell sequencing was performed following a previously published protocol *(33)* and manufacturer’s user guide (Document CG000186 Rev D, 10X Genomics). The complete protocol is available on protocols.io (https://www.protocols.io/view/10x-citeseq-protocol-covid-19-patient-samples-tetr-bqnqmvdw). In each experiment, PBMCs from 22-25 participants were used including 16-23 patients and 2-8 healthy individuals. Longitudinal samples of the same patient were used in different experiments to allow genetic demultiplexing. Each experiment used samples collected at different longitudinal time points to prevent that experimental conditions are aligning with potential batch effects.

In brief, PBMCs were thawed in a 37°C water bath for 30 s and washed with 5 mL of cRPMI followed by centrifugation at 350 rcf for 5 min at RT. Cell counts and viability were determined using a Vi-CELL XR Automated Cell Counter (Beckman Coulter Life Sciences). Equal number of cells were aliquoted from each sample to create a pool of 1.5×10^6^ cells with an average viability of 85% or higher. Pooled PBMCs were blocked with Human TruStain FcX (BioLegend) for 10 min on ice, followed by staining with a customized TotalSeq-C human cocktail for 45 min on ice (**Table S4**). Cells were washed three times, resuspended in PBS with 0.04% BSA, filtered through a 40 μm filter, and counted with Countess II Automated Cell Counter (Thermo Fisher Scientific).

Single cell suspensions were loaded into a Chromium Single Cell Chip A for single cell encapsulation using the 10X Chromium controller according to the manufacturer’s user guide (Document CG000186 Rev D, 10X Genomics), and as previously described *(34)*. In each experiment, the pooled cells of 22-25 participants were loaded into 4-6 individual lanes aiming for 7×10^4^ loaded cells per lane.

### Single-cell library preparation and sequencing

Single-cell libraries were constructed following the manufacturer’s user guide (Document CG000186 Rev D). cDNA libraries were generated using the Chromium Single-cell 5’ library & Gel bead kit and i7 Multiplex kit. Surface protein Feature Barcode libraries were generated with Chromium Single Cell 5′ Feature Barcode Library Kit and i7 Multiplex Kit N, Set A. In total, libraries generated from 971,550 cells were PE150 sequenced at the CZ Biohub on 18 lanes of an Illumina NovaSeq 6000 sequencer using a NovaSeq 6000 S4 Reagent Kit v1.

### Genotyping, sample demultiplexing and doublet detection

To assign cells to donors of origin in our multiplexed design, we used the genetic demultiplexing tools Freemuxlet and vcf-match-sample-ids, each a part of the Popscle suite of statistical genetics tools (https://github.com/statgen/popscle). Freemuxlet leverages the single-nucleotide polymorphisms (SNPs) present in transcripts and performs unsupervised clustering on the droplet barcodes to assign each to a nameless donor, or assign them as doublets between genetically-distinct nameless donors. The algorithm takes in a list of candidate loci throughout the genome at which to scan for SNPs, and returns droplet barcodes with donor assignments and a set of observed variants per donor. These sets of variants are then matched using genotypic similarity to those from an orthogonal bulk RNA-seq assay, done on an individual basis, to determine which donor is which patient. Once nameless donors are matched to uniquely-identifiable patients, droplet data are then joined with the other clinical covariates available for the patients, including age, sex, race, and disease status.

Freemuxlet was run on each of the 9 droplet reaction runs separately, using a list of exonic SNPs that were expected to be found in the 5’-end scRNA-seq data and that have a minor allele frequency > 0.05, based on data from the 1000 genomes project *(35)*.

### Bulk RNA-sequencing

Bulk RNA-seq data was generated to extract genotype information, so that single-cells could be demultiplexed and matched to single donors. For each donor, RNA was extracted from PBMCs using the Quick RNA MagBead kit (Zymo Research) on a KingFisher Flex system (Thermofisher Scientific) according to the company’s protocol. RNA integrity was measured with the Fragment Analyzer (Agilent) and library generation was continued when integrity was at least 6. Total RNA-sequencing libraries were depleted from ribosomal and hemoglobin RNAs, and generated using FastSelect (Qiagen) and Universal Plus mRNA-seq with Nu Quant (Tecan) reagents. Pooled libraries were PE100 sequenced on an HiSeq4000 or PE150 sequenced on an Illumina NovaSeq 6000 S4 flow cell at the CZ Biohub.

### Single-cell epitope and RNA-sequencing preprocessing and alignment

CellRanger v3.1.0 (run 1 to 7, cDNA library generation of run 6 failed) or v4.0.0 (run 8 to 10) software with the default settings was used to demultiplex the sequencing data and generate FASTQ files (Cellranger mkfastq), align the sequencing reads to the hg38 reference genome, and generate a unique molecular identifier (UMI)-filtered gene and protein expression count matrix for each lane (Cellranger count for scRNA-seq and CITE-seq data). Count matrices were then concatenated across all 50 lanes to generate two matrices: one mRNA matrix with 971,550 cells and 36,601 genes, and one surface protein matrix with 971,550 cells and 189 proteins.

### Single-cell epitope and RNA-sequencing processing and quality control

Resulting gene and protein expression count matrices were further processed in the Python package Scanpy v1.5.1 *(36)*. Processing of the concatenated mRNA count matrix was done using a novel two-step process. We have found empirically that traditional workflows for the processing of droplet-based RNA-seq data for PBMCs can sometimes create unwanted effects in the downstream endpoints. In particular, filtering of cells with a high percentage of mitochondrial cells may create visual artifacts in UMAP projections, and filtering of the matrix to only a few hundred highly-variable genes, while reducing the memory footprint of the data, can sometimes lead to spurious clustering of cells based on only a few genes. Our iterative process yields a UMAP projection that captures all available heterogeneity with minimal filtering in the first iteration, then removes non-target cells and corrects for non-biological signal in the second iteration. By doing this, we use a relatively large number of components to inform projections and clustering, but observe that the outputs in our dataset match the known biology better (e.g. proximity of similar cell types and states in UMAP space) and yield higher-confidence annotations.

In the first step, the mRNA matrix was filtered to remove doublet droplets, as annotated by freemuxlet, and very lowly-expressed genes with less than 100 UMIs across the 9 runs. The matrix was then normalized to yield a constant UMI sum per cell and log transformed. Matrix values were scaled to yield a mean of zero and standard deviation of 1, per gene. Principal component analysis (PCA) was performed, followed by nearest neighbors, UMAP projection and Leiden clustering, using an input of the 150 PCs with the highest variance explained and otherwise default Scanpy parameters. At this stage, Leiden clustering resolution was adjusted and restricted to certain clusters to separate out clusters of cells that projected into similar UMAP space. These clusters were subsequently annotated to mark those with a high percentage of mitochondrial content, which typically represent dead or dying cells, as well as mark clusters with high levels of hemoglobin and platelet factor expression, representing the non-target cell types of red blood cells and platelets. At this stage, we also observed that there were prominent batch effects in UMAP space that required correction.

In the second iteration, non-target cell types marked in the first step were removed prior to processing. Then, the same processing was followed starting from the raw data, with the exception that ComBat batch correction *(37)* was performed (to correct for the “run” covariate) after scaling and before performing PCA. Finally, further filtration of a relatively small number of cells with high expression of platelet, red blood cell, and mitochondrial genes was performed, as well as removal of donors that declined study participation. After processing, 600,929 cells and 18,262 genes remained in the mRNA matrix.

The surface protein matrix was filtered to the cells found in the mRNA matrix. One protein was removed from the matrix, as it appeared to have very low counts relative to the other surface proteins. The remaining proteins were then normalized using the centered-log ratio (CLR), computed for each gene independently. The CLR has typically been used for CITE-seq data with the recognition that antibody counts are typically not zero-inflated and FACS-like Gaussian distributions are achievable when treating the data as compositional. However, we recognized that the CLR-normalized distributions were affected by a relatively small number of cells that had extremely high or low expression, skewing the visualization of the Gaussian mixture distributions. To remedy this effect, we identified boundary values of the distributions for each gene using a bin height threshold when values were plotted on a histogram, clipped the values to these boundaries, and scaled the remaining values between 0 and 1.

### Cell type classification

After processing, Leiden clustering was adjusted to match the clustering of cells projected into UMAP space. Cell type annotation was performed at 3 levels of granularity based on known marker gene and protein expression, as well as differentially expressed genes between clusters using a Wilcoxon rank-sum test. At the lowest level of granularity, we identified 11 cell types corresponding to the known major cell types present in PBMCs: T4, T8, Tgd, cMs, ncMs, NK, B, PBs, cDC, pDC and Progen cells. At the next level of granularity, we separate out memory, naïve, and proliferating subtypes in the lymphocytes; two different subtypes known in each of the NK cells, and conventional DCs; regulatory T cells (Tregs) from the T4 group; mucosal-associated invariant T (MAIT) cells from the T8 group; subpopulations of lineage-committed progenitor cells; and a subpopulation of B cells that seemed to be committed to the PB lineage. At the highest level of granularity, we further separate out a CD8+ effector memory population, an NK population with CD3 transcript expression, early and late proliferating subpopulations in the lymphocytes, and a few subpopulations that were either donor-specific (patients 1002 in the B cells and 1006 in the monocytes) or run-specific (i.e. cells from run 3, which exhibited some processing issues and for which ComBat *(37)* was unable to correct).

### Differential proportion analysis

Differences in cell type proportions were assessed in C19+ versus C19- cases or Healthy controls by aggregating cell type observations per COVID status at timepoint D0. Additionally, in the C19+ cases for which all 4 timepoints were available, cell type proportion changes were assessed over time. Differential proportion analysis was performed using a permutation-based approach that compares observed cell type proportion differences with those calculated from a null-distribution that is generated by randomly shuffling cell type labels (100,000 permutations) for a fraction (w=0.1) of the total cells, as described previously *(38)*. Resulting p-values were corrected for multiple testing using Holm’s correction, after which an adjusted p-value of <0.05 was considered significant.

### Differential expression analysis

Differences in gene expression levels were determined for each of the myleoid cell types between C19+ cases at D0, D4, D7 or D14 versus Healthy controls. To assess whether these changes are specific for C19+ cases or are a more general phenomenon as a consequence of acute respiratory distress syndrome, we also compared the upregulated genes with the C19- cases. Differential expression analysis was performed per run on the raw gene expression matrix using Memento v0.0.4 (unpublished, Kim MC et al. https://github.com/yelabucsf/scrna-parameter-estimation), after which results were meta-analyzed over all runs. Genes were pre-filtered based on a minimum raw mean expression of 0.07 within at least 90% of both comparison group. A false discovery rate of <0.05 was used to determine statistical significance.

### Differential expression heatmaps

Heatmaps show the pseudobulked, Z-scored expression values of the donors present at each time point for the top significantly upregulated genes. To generate the heatmaps, cells were first subsetted from the larger mRNA matrix to only those of a given cell type. Counts were pseudobulked across all genes by patient present at each time point, yielding a single gene-by-sample matrix, with 179 unique donor-timepoint samples. The genes in this matrix were subsetted to the union of the top 150 genes with the highest differential expression coefficient at each timepoint, using the 1-dimensional memento results that tested gene counts in C19+ cases vs. healthy controls. Genes were further filtered to remove those that had high variance in healthy controls (standard deviation > 0.5), since these were enriched for what seemed to be a non-biological signal (e.g. ribosome-associated genes). The matrix, now with 204 genes, was then Z scored and separated by time point to 4 matrices, with healthy samples being distributed to each matrix.

Ordering of the rows and columns were computed such that they would be consistent among the heat maps. Genes were clustered by k means using only the values of the day 0 time point, with *k*=6 chosen by the “elbow” point of the graph plotting distortion (using a sum of square errors cost function) with increasing numbers of clusters. Columns of each heatmap were determined by taking the 80 columns across all heatmaps that had the earliest time point for each donor, subsetting according to their disease status (Healthy, COVID-19 negative, and COVID-19 positive), and then hierarchically clustering within each of those groups. With this ordering, each donor then had a unique position along the horizontal axis, which was then applied to all the heatmaps, omitting those samples that were absent from a given time point. GSEA was done using the GOATOOLS Python package *(39)*, filtering to terms with at least 2 associated genes.

### Interferon Stimulated Gene Score Method

An orthogonal scRNA-seq dataset containing PBMCs stimulated with IFN beta and gamma was used to identify the specific and shared type I and type II ISGs in the cMs (unpublished). The gene list was used to calculate a type I, type II and shared ISG score based on the average gene expression count of the unique or shared type I and type II ISGs. These ISG scores were calculated for each unique combination of cell type, donor and timepoint. Subsequently, ISG scores were averaged over each of the disease categories (C19+ moderate/severe, C19- moderate/severe, healthy control) and then log2-transformed. A Welch’s T-test was performed to compare the ISG score between C19+ patients and healthy control. Significance was defined as Bonferroni-adjusted p-value < 0.05.

### SARS-CoV-2 detection by clinical qRT-PCR

Viral titers were quantified in a subset of the C19+ patients in the UCSF CLIAHUB Clinical Microbiology Laboratory. For this, RNA was extracted from tracheal aspirate samples and used for qRT-PCR as previously described *(40)*. In short, viral titers were assessed using primers targeting the SARS-CoV-2 N gene (Ct1), E gene (Ct2) and human RNAse P gene (Ct_host, positive control). The Ct value of the viral N or E gene was subtracted from the human RNAse P gene (delta Ct: dCt1 and dCt2) and number signs were reversed to obtain a measurement for viral load. As there was an almost perfect correlation between dCt1 and dCt2 values (Pearson R = 0.99, p = 1.9 x 10^−283^) and dCt2 had the least missing values, viral load is represented by the dCt2 values. dCt2 values as measured in the tracheal aspirate samples were linked to the scRNA-seq PBMC data of the same donor, and the closest possible timepoint (up to 2 days apart).

### Radioligand binding assay for anti-IFN-α2 autoantibody detection

A DNA plasmid containing full-length cDNA sequence with a Flag-Myc tag (Origene #RC221091) was verified by Sanger sequencing and used as template in T7-promoter-based *in vitro* transcription/translation reactions (Promega, Madison, WI: #L1170) using [S35]-methionine (PerkinELmer, Waltham, MA; #NEG709A). IFN-α2 protein was column-purified using Nap-5 columns (GE Healthcare, Chicago, IL; #17-0853-01), incubated with 2.5ul serum, or 2.5ul plasma, or 1ul anti-myc positive control antibody (CellSignal, Danvers, MA; #2272), and immunoprecipitated with Sephadex protein A/G beads (Sigma Aldrich, St. Louis, MO; #GE17-5280-02 and #GE17-0618-05, 4:1 ratio) in 96-well polyvinylidene difluoride filtration plates (Corning, Corning, NY; #EK-680860). The radioactive counts (cpms) of immunoprecipitated protein was quantified using a 96-well Microbeta Trilux liquid scintillation plate reader (Perkin Elmer). Antibody index for each sample was calculated as follows: (sample cpm value – mean blank cpm value) / (positive control antibody cpm value – mean blank cpm value). For the COVID-19 patient and convalescent plasma cohorts, a positive signal was defined as greater than 6 standard deviations above the mean of pre-COVID-19 blood bank non-inflammatory controls. For the large asymptomatic San Francisco community population cohort, a positive signal was defined as having a z-score greater than 3.3 (p=0.0005) relative to the whole cohort.

### Luciferase reporter assays

The blocking activity of anti-IFN-α autoantibodies was determined by assessing a reporter luciferase activity. Briefly, HEK293T cells were transfected with the firefly luciferase plasmids under the control human *ISRE* promoters in the pGL4.45 backbone, and a constitutively expressing *Renilla* luciferase plasmid for normalization (pRL-SV40). Cells were transfected in the presence of the X-tremeGene 9 transfection reagent (Sigma Aldrich) for 36 hours. The, Dulbecco’s modified Eagle medium (DMEM, Thermo Fisher Scientific) medium supplemented with 10% healthy control or patient serum/plasma and were either left unstimulated or were stimulated with IFN-α, IFN-ω or IFN-β (10 ng/mL) for 16 hours at 37°C. Each sample was tested once. Finally, Luciferase levels were measured with the Dual-Glo reagent, according to the manufacturer’s protocol (Promega). Firefly luciferase values were normalized against *Renilla* luciferase values, and fold induction is calculated relative to controls transfected with empty plasmids.

## Data availability

Processed (deanonymized) single-cell RNA-sequencing data has been deposited in the Gene Expression Omnibus under the accession number GSE168453 and is currently being deposited in the data coordinate platform (DCP) of the Chan Zuckerberg Initiative (CZI) Human Cell Atlas.

## Code availability

The original Python code for Scanpy (https://github.com/theislab/scanpy), Freemuxlet ((https://github.com/statgen/popscle), can be found at Github. All custom-made code is currently being checked into GitHub repository (https://github.com/yelabucsf/COVID-19).

## Acknowlegements

We thank the patients and their families for placing their trust in us. We thank all members of the Ye, Anderson, DeRisi and Casanova Labs for helpful discussions. This study was performed with support from the National Institute of Allergy and Infectious Diseases-sponsored Immunophenotyping Assessment in a COVID-19 Cohort (IMPACC) Network (National Institute of Allergy and Infectious Diseases grant U19 AI1077439 to D.J.E). This work was supported by grants from the Dutch Research Council (NWO-Veni 192.029 to M.G.P.W.), the National Institute of Diabetes, Digestive and Kidney Diseases (1F30DK123915-01 to S.E.V.), the Chan Zuckerberg Biohub, the National Institute of Allergy and Infectious Diseases (5PO1AI118688-04 to M.S.A) and the National Heart, Lung and Blood Institute (R35 HL140026 to C.S.C.). AleW and Y.S.S. were supported in part by an NIH grant R35-GM134922 and the Exascale Computing Project (17-SC-20-SC), a collaborative effort of the U.S. Department of Energy Office of Science and the National Nuclear Security Administration. The Laboratory of Human Genetics of Infectious Diseases is supported by the Howard Hughes Medical Institute, the Rockefeller University, the St. Giles Foundation, the National Institutes of Health (NIH) (R01AI088364), the National Center for Advancing Translational Sciences (NCATS), NIH Clinical and Translational Science Award (CTSA) program (UL1 TR001866), a Fast Grant from Emergent Ventures, Mercatus Center at George Mason University, the Yale Center for Mendelian Genomics and the GSP Coordinating Center funded by the National Human Genome Research Institute (NHGRI) (UM1HG006504 and U24HG008956), the French National Research Agency (ANR) under the “Investments for the Future” program (ANR-10-IAHU-01), the Integrative Biology of Emerging Infectious Diseases Laboratory of Excellence (ANR-10-LABX-62-IBEID), the French Foundation for Medical Research (FRM) (EQU201903007798), the FRM and ANR GENCOVID project, ANRS-COV05, the Square Foundation, *Grandir - Fonds de solidarité pour l’enfance*, the SCOR Corporate Foundation for Science, Institut National de la Santé et de la Recherche Médicale (INSERM) and the University of Paris. The work was supported in part by the Intramural Research Program of the NIAID, NIH. P.B. and T.L.V. were supported by the MD-PhD program of the Imagine Institute (with the support of the Fondation Bettencourt-Schueller).

