## Supplementary Figure S1-S3 for "Longitudinal single-cell epitope and RNA-sequencing reveals the immunological impact of type 1 interferon autoantibodies in critical COVID-19"

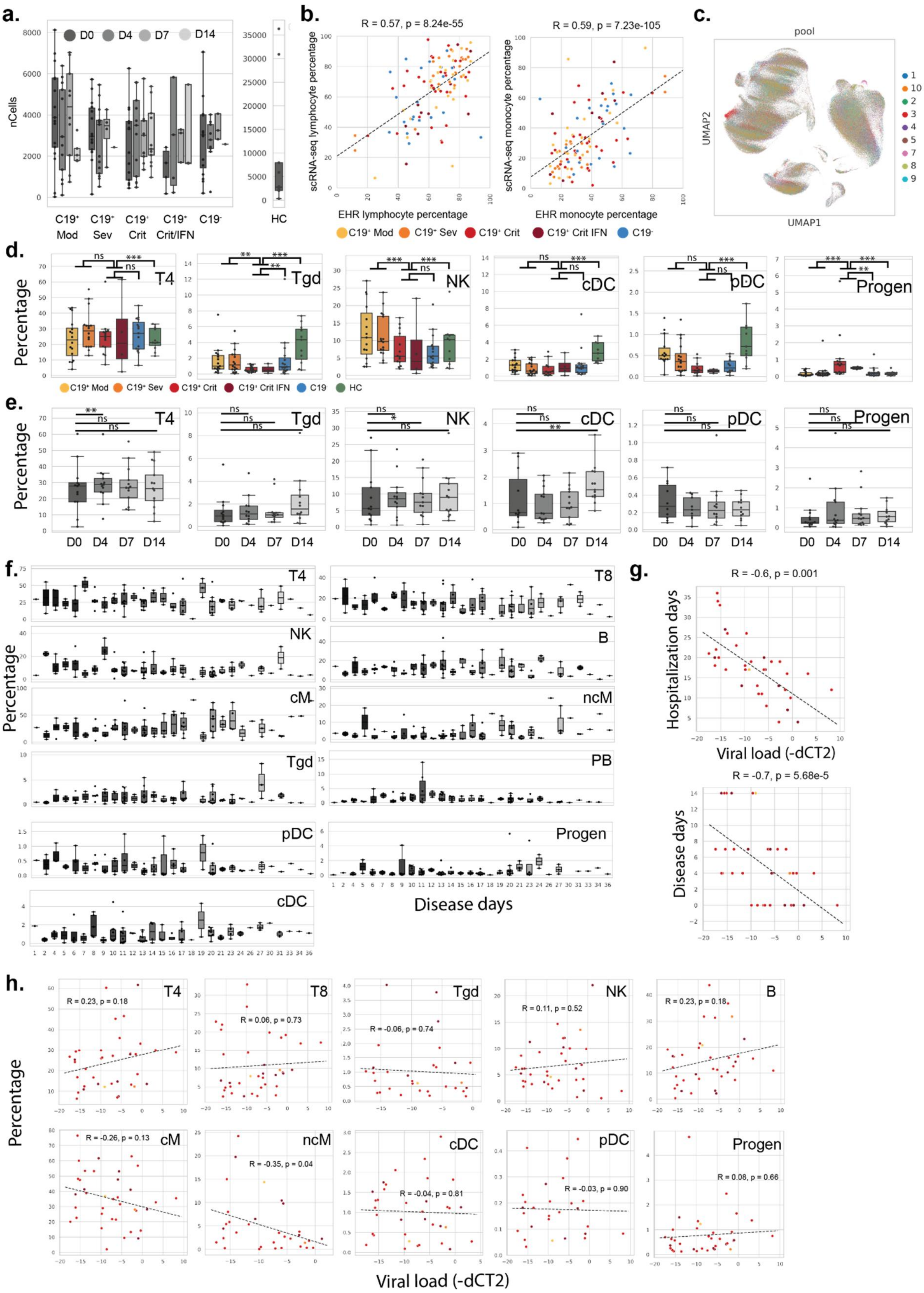

a.
D0 D4 D7 D14
nCells
C19+ Mod
C19+ Sev
C19+ Crit
C19+ Crit/IFN
C19-
HC
b.
R = 0.57, p = 8.24e-55
scRNA-seq lymphocyte percentage
EHR lymphocyte percentage
R = 0.59, p = 7.23e-105
scRNA-seq monocyte percentage
EHR monocyte percentage
C19+ Mod C19+ Sev C19+ Crit C19+ Crit IFN C19-
c.
pool
UMAP2
UMAP1
d.
Percentage
T4
Tgd
NK
cDC
pDC
Progen
C19+ Mod C19+ Sev C19+ Crit C19+ Crit IFN C19 HC
e.
Percentage
T4
Tgd
NK
cDC
pDC
Progen
D0 D4 D7 D14
f.
Percentage
T4
T8
NK
B
cM
ncM
Tgd
PB
pDC
Progen
cDC
Disease days
g.
R = -0.6, p = 0.001
Hospitalization days
Viral load (-dCT2)
R = -0.7, p = 5.68e-5
Disease days
h.
Percentage
T4 R = 0.23, p = 0.18
T8 R = 0.06, p = 0.73
Tgd R = -0.06, p = 0.74
NK R = 0.11, p = 0.52
B R = 0.23, p = 0.18
cM R = -0.26, p = 0.13
ncM R = -0.35, p = 0.04
cDC R = -0.04, p = 0.81
pDC R = -0.03, p = 0.90
Progen R = 0.08, p = 0.66
Viral load (-dCT2)

**Figure S1: Shifts in circulating leukocyte composition and correlations with SARS-CoV2 titers in critical COVID-19. a)** Average number of cells remaining after QC per donor per timepoint per disease severity status. **b)** Pearson correlation between electronic health record (EHR) and scRNA-seq derived blood counts for lymphocytes (left) and monocytes (right). Each datapoint is colored by disease severity status. **c)** UMAP projection of PBMCs colored by experimental run. **d)** Boxplots (showing median, $25^{th}$ and $75^{th}$ percentile) of the percentages of T4, Tgd, NK, cDC, pDC, Progen cells (y-axis) by COVID-19 status and severity level on day of hospital admission (D0). Each dot represents the percentage of a specific cell type per donor. Shown statistical comparisons are between cells from all C19+ critical donors (including the anti-IFN-$\alpha$2 autoantibody donors) and healthy controls (HC), C19- donors or combined C19+ Moderate-Severe donors. **e)** Boxplots of the percentages of T4, Tgd, NK, cDC, pDC, Progen cells (y-axis) in COVID-19 patients over day 0, 4, 7 and 14 since hospitalization (D0, D4, D7, D14). **f)** Boxplots of the percentages of T4, T8, Tgd, NK, B, PB, cM, ncM, cDC, pDC, Progen cells (y-axis) by COVID-19 status and severity level on day since first symptoms. **g)** Scatterplot of SARS-CoV2 viral titer as measured by qRT-PCR in tracheal aspirates (inverse dCT, x-axis) and days since hospitalization (left) or days since first symptoms (rights) (R = Pearson correlation). **h)** Scatterplot of SARS-CoV2 viral titer and cell type compositions of total PBMCs of T4, T8, Tgd, NK, B, PB, cM, ncM, cDC, pDC, Progen cells (R = Pearson correlation). *** $p < 0.001$, ** $p < 0.01$, * $p < 0.05$, ns = not significant.

#### Figure S2

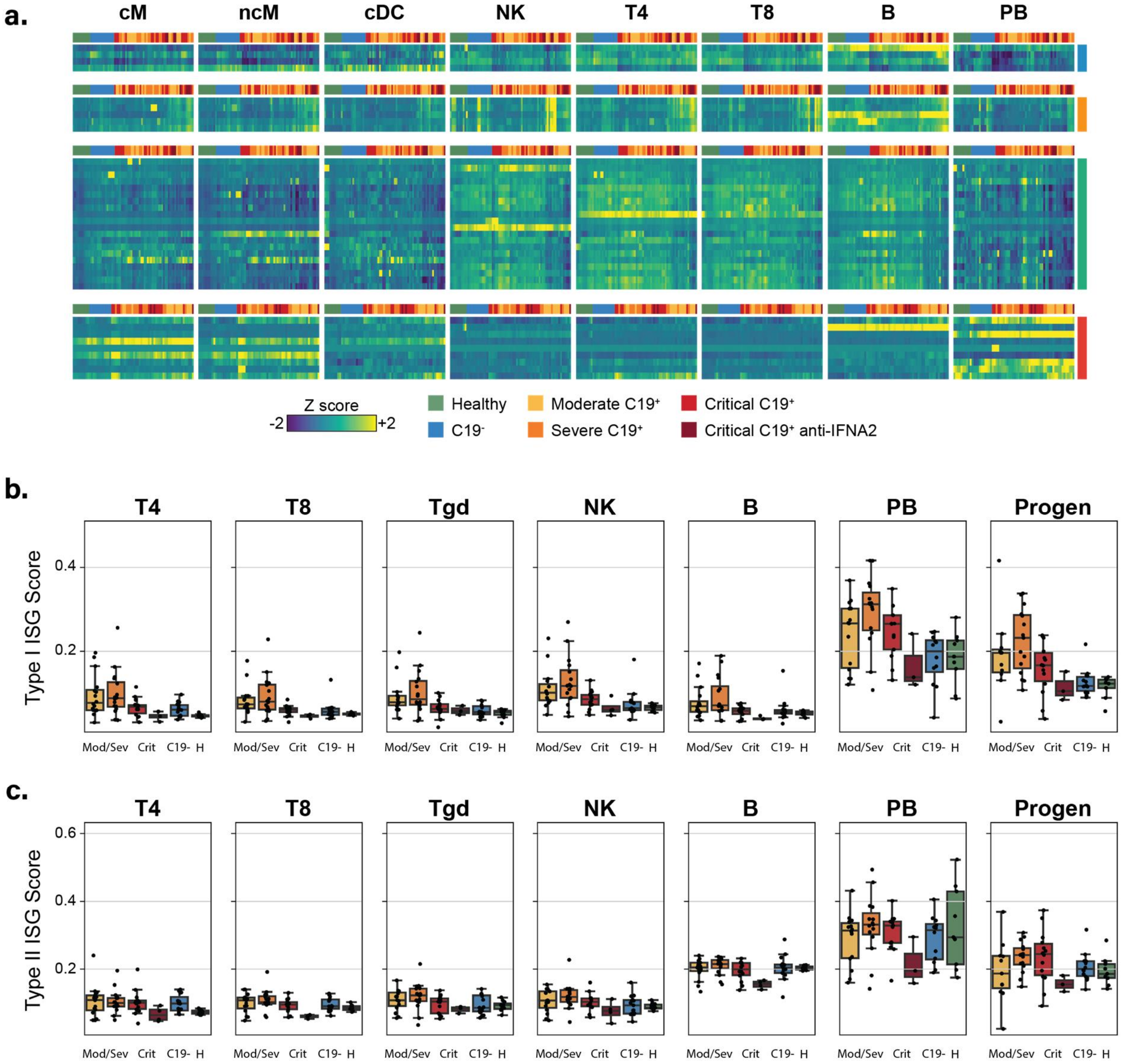


**Figure S2: Feature abundance changes of leukocyte subsets in critical COVID19. a)** Heatmap of 38 surface proteins differentially expressed at day 0 (FDR < 0.05, |log(fold change)| >0) in at least one of the 11 cell types. CD4+ T cells (T4), CD8+ T cells (T8), natural killer cells (NK), B cells (B), plasmablasts (PB), classical monocytes (cM), non-classical monocytes (ncM), and conventional dendritic cells (cDC) are shown. Each row represents a surface protein and each column is the average expression of the proteins in a particular sample across all cells of a specific type. Samples are grouped by case-control status and C19+ severity. Expression levels are row standardized. **b)** Type I-specific ISG score (y-axis) at day 0 across 8 cell types separated by case-control status and disease severity. **c)** Type II-specific ISG score (y-axis) at day 0 across 8 cell types separated by case-control status and disease severity. Boxplots show median, 25$^{th}$ and 75$^{th}$ percentile.

### Figure S3

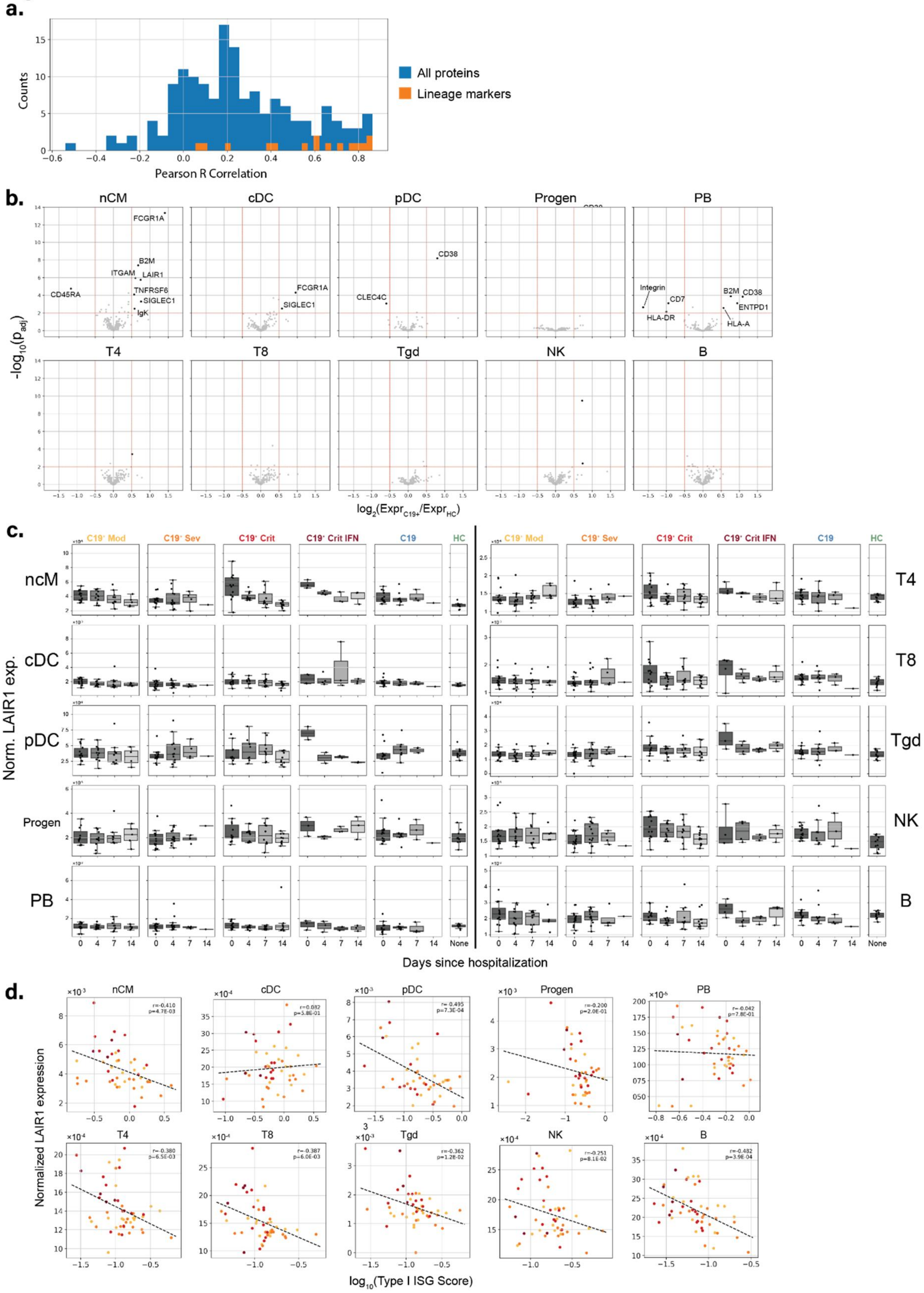
a.
Counts
Pearson R Correlation
All proteins
Lineage markers
b.
nCM
cDC
pDC
Progen
PB
T4
T8
Tgd
NK
B
FCGR1A
B2M
ITGAM
LAIR1
TNFRSF6
SIGLEC1
IgK
CD45RA
CLEC4C
CD38
Integrin
CD7
HLA-DR
ENTPD1
HLA-A
-log10(padj)
log2(ExprC19+/ExprHC)
c.
C19+ Mod
C19+ Sev
C19+ Crit
C19+ Crit IFN
C19
HC
Norm. LAIR1 exp.
Days since hospitalization
None
d.
Normalized LAIR1 expression
log10(Type I ISG Score)

**Figure S3: Surface protein abundance changes of leukocyte subsets in critical COVID19. a)** Density plot of Pearson R correlations between normalized protein expression and corresponding transcript expression for each cell type in each sample. Correlations for 15 lineage markers highlighted in orange. **b)** Volcano plot of log fold change between C19+ and healthy controls (x-axis) versus -log10(P-value) (y-axis) for 10 additional cell types. Proteins that are statistically significant (FDR < 0.05) and have a log2(fold change) > 0.5 are highlighted. **c)** Normalized LAIR-1 surface expression (y-axis) for each cell type over the course of disease for healthy controls, C19- controls, and C19+ cases. C19+ cases are separated by disease severity and the presence of anti-IFN-$\alpha$2 antibodies. **d)** Scatterplot of normalized LAIR-1 expression (y-axis) versus the type I-specific ISG score (x-axis) for additional cell types, showing C19+ cases colored by severity and anti-IFN-$\alpha$2 status.
